# Hemisphere-Specific Properties of the Ventriloquism Aftereffect in Humans and Monkeys

**DOI:** 10.1101/564682

**Authors:** Norbert Kopčo, Peter Lokša, I-fan Lin, Jennifer Groh, Barbara Shinn-Cunningham

## Abstract

Visual calibration of auditory space requires re-alignment of representations differing in 1) format (auditory hemispheric channels vs. visual maps) and 2) reference frames (head-centered vs. eye-centered). Here, a ventriloquism paradigm from Kopčo *et al*. (J Neurosci, 29, 13809-13814) was used to examine these processes in humans and monkeys for ventriloquism induced within one spatial hemifield. Results show that 1) the auditory representation is adapted even by aligned audio-visual stimuli, and 2) the spatial reference frame is primarily head-centered in humans but mixed in monkeys. These results support the view that the ventriloquism aftereffect is driven by multiple spatially non-uniform processes.

PACS numbers: 43.66.Pn, 43.66.Qp, 43.66.Mk

## 1. Introduction

Vision plays an important role in calibration of auditory spatial perception. In the “ventriloquism aftereffect” (VAE), repeated presentations of spatially mismatched visual and auditory stimuli produce a shift in perceived sound location that persists when the sound is presented alone (Canon, 1970; Recanzone, 1998; Woods and Recanzone, 2004; Bertelson *et al.*, 2006). The brain mechanisms that support this process are mysterious because spatial representations seem to differ in vision and in hearing in two ways.

First, visual space is initially encoded relative to the direction of the eye gaze, while the cues for auditory space are first computed relative to the orientation of the head (Groh & Sparks, 1992). A means of reconciling this discrepancy in reference frames (RF) is necessary to achieve correct recalibration. Our previous study suggests that a mixture of eye-centered and head-centered RFs are associated with recalibration in the central region of the audiovisual field (Kopco *et al.*, 2009).

Second, there is growing evidence that, in mammals, auditory space is encoded non-homogeneously, based on two (or more) spatial channels roughly aligned with the left and right hemifields of the horizontal plane (Grothe *et al.*, 2010; Groh, 2014). This is markedly different from visual spatial codes, in which the retinal surface provides a map of the position of stimuli in the environment.

Thus, the process of using visual information to recalibrate auditory space is multifaceted, and may operate differently in different portions of the environmental scene. Indeed, differential patterns of adaptation across auditory space have been observed (Phillips and Hall, 2005; Maier *et al.*, 2010), suggesting that the auditory code in humans likely employs the same two-channel scheme that has been observed in animal species (Salminen *et al.*, 2009).

Here, we tested whether the spatial characteristics of the ventriloquism aftereffect induced in the audiovisual periphery (i.e., in a single hemifield) differ from those occurring when the aftereffect is induced in the central region (i.e., covering both hemifields; (Kopco *et al.*, 2009)). Persistent visually driven biases in perceived sound location were induced in seven humans and two monkeys. As in Kopco et al. (2009), we presented the mismatched (5-6°-shifted) audio-visual (AV) stimuli in only a subregion of space (Fig. 1A, top panel), but this time the training region was peripheral, rather than central, to the fixation point used for these trials (Fig. 1A, bottom panel). We evaluated the effects of this pairing on saccade accuracy for interleaved auditory-only trials both from that fixation point and a non-training fixation point in the opposite hemifield (Fig. 1A, bottom panel).

**Fig. 1.**
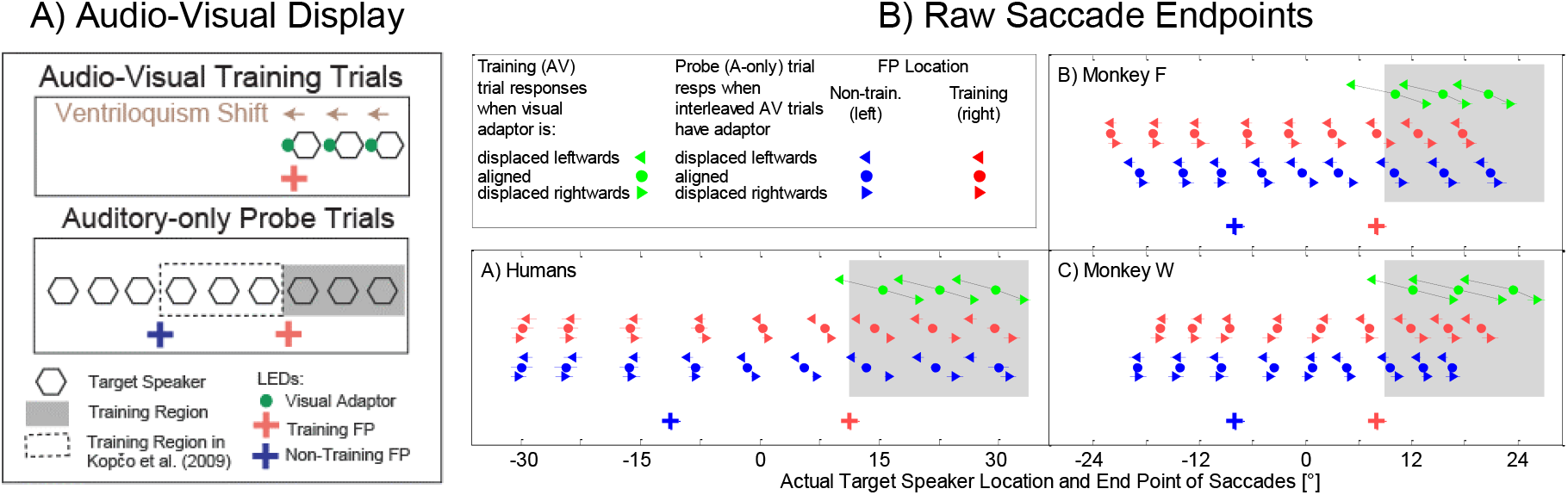
Experimental set-up and raw experimental data. A) Audiovisual display used to present the AV training stimuli in one experimental block. At the beginning of each AV training trial (top), the subject had to fixate on the same initial FP; then, the training stimulus was presented from one of three locations lateral to the FP, keeping the direction of the induced shift the same within a block (by consistently presenting the visual adaptor displaced to the left, to the right, or aligned with the target speaker). On the auditory-only probe trials (bottom), the same nine speaker locations and two FPs were used in all blocks. The probe trials were randomly interleaved among the training trials and the FP and target locations varied randomly from trial to trial. Dashed frame indicates the central training region used in Kopco et al. (2009). B) Raw saccade endpoints of the responses to the AV training stimuli and auditory-only probe stimuli as a function of the actual target speaker location, collapsed across time. The symbols represent responses in different audiovisual conditions (see legend), separately for the training trials (green), probe trials starting at the training fixation (red), and probe trials starting at the nontraining fixation (blue). Left panel: Across-human-subject mean (+/-1 SEM) of responses. Right panels: Individual monkeys’ across-block means (+/-1 SEM) of responses. The dashed lines connect symbol triplets for the same auditory target when presented with one of the three different visual adaptors (symbol triplets for A-only responses corresponding to the same target location are not explicitly connected as they are not confusable). Graphs for each measurement type are plotted in one row, vertically offset from data for other types, for visual clarity.

As was the case for our previous study involving central training, the pairing of a displaced visual stimulus induced an aftereffect in the trained region. In monkeys, the reference frame was similar to our previous findings (a hybrid of head- and eye-centered coordinates), whereas in humans, the contribution of an eye-centered component was not readily apparent. However, in both species, we also observed biases related to the location of the fixation point, even when the AV stimuli were aligned. Together, these findings confirm the contribution of multiple signals related to different reference frames and representational formats across the horizontal space.

## 2. Methods

All procedures and equipment closely matched those used in Kopco *et al.* (2009).

### General methods

Subjects made eye movements from a visual fixation point to a broadband noise delivered from loudspeakers in darkness. On training trials (Fig. 1A, top), visual stimuli were presented simultaneously with the sounds, using light-emitting diodes (LEDs) displaced from the locations of the speakers. On randomly interleaved probe trials (Fig. 1A, bottom), only the auditory stimuli were presented (50% of all trials).

### Subjects

Seven human subjects and two adult male rhesus monkeys participated. The human and animal experimental protocols were approved by the institutional review committees at Boston University and Duke University, respectively.

### Setup

Subjects were seated in a quiet darkened room in front of an array of speakers and LEDs (Fig. 1). To keep the head-centered RF fixed, the subjects’ heads were restrained (humans, chin rest; monkeys, implanted head post). Subjects’ behavior was monitored and responses were collected by an infrared eye tracker (humans) or implanted scleral eye coil (monkeys). The eye-tracking system was calibrated using visually guided saccades to selected target locations at the beginning of each session.

### Stimuli

Sounds were broadband noises with 10 ms on/off ramps [humans, 100 ms, 0.2–6 kHz, 70 dBA; monkeys, ~500–1000 ms, 0.5–18 kHz, 50 dBA] presented from speakers mounted on the horizontal plane ~1.2 m (humans) or 1.45 m (monkeys) from the center of the listener’s head. Spacing between speakers was 7.5° (humans) or 6° (monkeys). The LEDs for the AV stimuli were mounted so that they were either horizontally aligned with the speakers or displaced (either to the left or to the right) by 5° (humans) or 6° (monkeys). They were turned on and off in synchrony with the corresponding speakers. Two additional LEDs 10° (humans) or 8° (monkeys) below the speaker array served as fixation locations (azimuths of ±11.8° in humans, ±8° in monkeys).

### Procedures

Trials began with the onset of one of the two fixation LEDs. After subjects fixated the LED for 150 ms (humans) or 500 ms (monkeys), the fixation LED was turned off and the AV or A-only stimulus was presented. The subjects performed a saccade to the perceived location of the stimulus (humans were instructed to look to the location of the auditory component of the stimulus; monkeys were rewarded for a saccade that ended within a 16°-wide rectangular window centered on the auditory component and covering the visual component on the AV trials).

Training (AV) and probe (A-only) trials were randomly interleaved at a ratio of 1:1 (in the monkeys, 6.75% of the total trials were AV-aligned and presented from the ±30° locations, just outside the range of the A-only test trials, to reduce the compressive bias observed even in the baseline responses; see Fig. 2). Trials were run in blocks with a consistent AV pairing (leftward, rightward, or no shift). For the monkeys, multiple blocks were conducted per session, with shifts in a particular direction for that session interleaved with no-shift blocks. For the humans, each session contained only one block and the order of blocks was random across the subjects. Each monkey performed a total of 128–160 blocks of ~600 trials each. Each human performed 12 sessions of ~720 trials each.

**Fig. 2.**
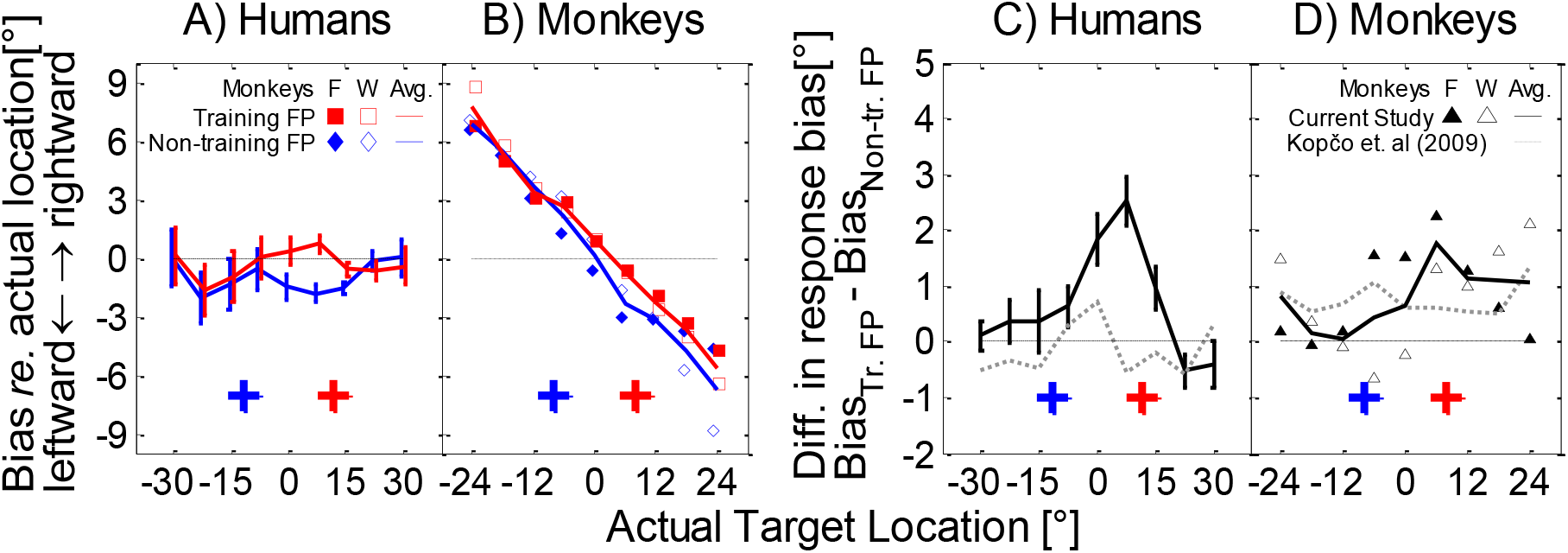
Average bias in A-only responses in the baseline conditions in which no AV shift was included plotted as a function of the actual target location (Panels A and C – humans; panels B and D – monkeys; filled symbols – monkey F, open symbols – monkey W)). The symbols at the bottom of the figures show the training fixation point location (red plus) and the non-training fixation point location (blue plus). Panels A and B show mean response biases when eyes are fixated at the training FP (red) and the non-training FP (blue). Panels C and D show the difference between responses from training FP and the non-training FP (difference between red and blue lines from the respective panels above) – the black solid lines. The gray dotted lines show the difference for central adaptation, taken from Kopco et al. (2009), for comparison purposes.

### Data analysis

Data from the first quarter of each block were excluded to remove transitory values observed during the initial buildup of VAE. Within-block averages were computed from the remaining data separately for each combination of target location, fixation position, and condition. Since no large left-right differences were observed, data with training FP on the left were mirror-flipped and combined with the data with training FP on the right. All human data are presented as across-subject means and standard errors of the mean, and all monkey data are presented as across-monkey means and the individual monkey data points.

## 3. Overall Design and Results

As in Kopco et al. (2009), we presented paired visual-auditory stimuli in a subregion of audiovisual space, fixed in both eye- and head-centered coordinates. We used one initial eye fixation position on training trials and presented the discrepant audiovisual stimuli from a restricted spatial range that was lateral with respect to the fixation point (see Fig. 1A, top). Because the visual training was local, we could test the spatial attributes of the resulting recalibration by shifting fixation on probe trials. Specifically, on interleaved auditory-only probe trials, we varied initial eye position with respect to the head (which was fixed) and presented sounds from all target locations spanning both the same head-centered locations and the same eye-centered locations as on the training trials (see Fig. 1A, bottom). We first consider the effects observed on the AV training trials themselves before turning to aspects of how the effects generalize to the auditory-only conditions across both the trained and untrained regions of space as a function of eye-referenced location, head-referenced location, and fixation position.

### 3.1 Ventriloquism effect

A strong ventriloquism *effect* – or capture of the auditory stimulus location by the visual stimulus on combined AV trials - was observed in both the humans and the monkeys. Green symbols in Fig. 1B show the raw responses in the humans (left-hand panel) and in the monkeys (right-hand panels). When the AV stimuli were aligned, the average responses were not biased at all in humans and medially biased by 1.5° in monkeys. The relative strength of the ventriloquism effect, expressed as percent of shift in responses towards the visual (V) component re. the A-component on misaligned AV trials, ranged from 82% to 96% in humans, and from 58% to 73% in monkeys (averaged across 2 directions of induced shift). Even though these results show that there was a decrease in the strength of the ventriloquism effect for the most lateral targets, it was expected that, as in Kopco *et al.* (2009), this strong ventriloquism *effect* would result in a clear local ventriloquism *after*effect.

### 3.2 Gaze-dependent Effects During AV-aligned Baseline

We next assessed the auditory-only responses interleaved with the spatially aligned AV stimuli. The red and blue circles in Fig. 1B show these responses. Overall, the pattern of results shows that both humans and monkeys accurately localized the auditory targets, showing a systematic displacement of the responses with the actual target locations. To analyze the impact of the visual training in more detail, the left-hand panels of Fig. 2 show the biases in responses to the A-only stimuli in the baseline condition for the humans (panel A) and monkeys (panel B). One visible difference between the species is that the monkeys showed a strong compressive bias (towards the center) in their responses. Humans’ responses showed no such effect. As discussed in Kopco *et al.* (2009), this difference is most likely due to different strategies the monkeys use in their responses, often making two saccades to reach an auditory target (Jay and Sparks, 1990). However, given that the current study analyzed the relative effect of the positive and negative VAE with respect to the baseline, such compressive biases are not expected to affect the results other than making the effects appear smaller.

The gaze-direction-dependent adaptation is seen when comparing the responses from the training FP (red) to those from the non-training FP (blue). In both species, the responses to the targets at approximately 10° azimuth were biased to the right by 2-3° when performed from the training FP (red “+” symbol) compared to the responses from the non-training FP (blue “+” symbol). Solid lines in panels C and D show the differences between the red and blue lines from panels A and B, respectively, while dotted lines represent the same data from the central-adaptation experiment of Kopco *et al.* (2009). These panels show that responses to auditory-only stimuli from AV-trained locations that are lateral and near the training FP differ depending on whether eyes fixate within the same hemifield or the opposite hemifield. On the other hand, when the AV training locations are in the center, covering both hemifields, no such differential effect of fixation location is observed (dashed lines). ANOVAs performed on the data from panels C and D showed a significant effect of target location for the humans (F_8,48_ = 9.45, p < 0.00001), but not for the monkeys (F_8,8_ = 0.96, p > 0.1), even though there is a clear trend for both monkeys to show auditory-only response bias at the locations around 10°. This effect of eye fixation direction is strong, of size comparable to the VAE (see next section), and it demonstrates that there is some eye-gaze-dependent contribution to responses to auditory-only stimuli even when vision is not used to induce any recalibration of the auditory spatial representation. However, this contribution is only visible if the AV stimuli are presented within one spatial hemifield.

### 3.3 Ventriloquism Aftereffect and its Reference Frame

The expected pattern of ventriloquism aftereffect, and the predictions about the reference frame based on it, are illustrated in Fig. 3A. The red line in the upper panel shows the predicted magnitude of the aftereffect induced by the AV stimuli, peaking in the trained region (gray region) when assessed with eyes fixating the training FP. If visually induced spatial plasticity occurs in a brain area using a head-centered RF, then shifts in perceived sound location should occur mainly for sounds at the same head-centered locations (in Fig. 3A, solid blue line matches the red line). Conversely, if plasticity occurs in an eye-centered RF, then visually induced shifts should occur mainly for sounds at the same eye-centered locations (dotted blue line is shifted to the left of the red line by the same displacement as the non-training FP is shifted relative to the training FP). The bottom panel of Figure 3A summarizes the predicted results if evaluated as a difference between the responses from the training and non-training FPs. The solid orange line shows the difference between the red and solid blue lines, corresponding to the expected results if the reference frame is head-centered. The dashed orange line shows the difference between the red and dashed blue lines, corresponding to the expected results if the reference frame is eye-centered. This panel also shows the difference in the biases expected if the RF is mixed, as observed in Kopco *et al.* (2009).

**Fig. 3.**
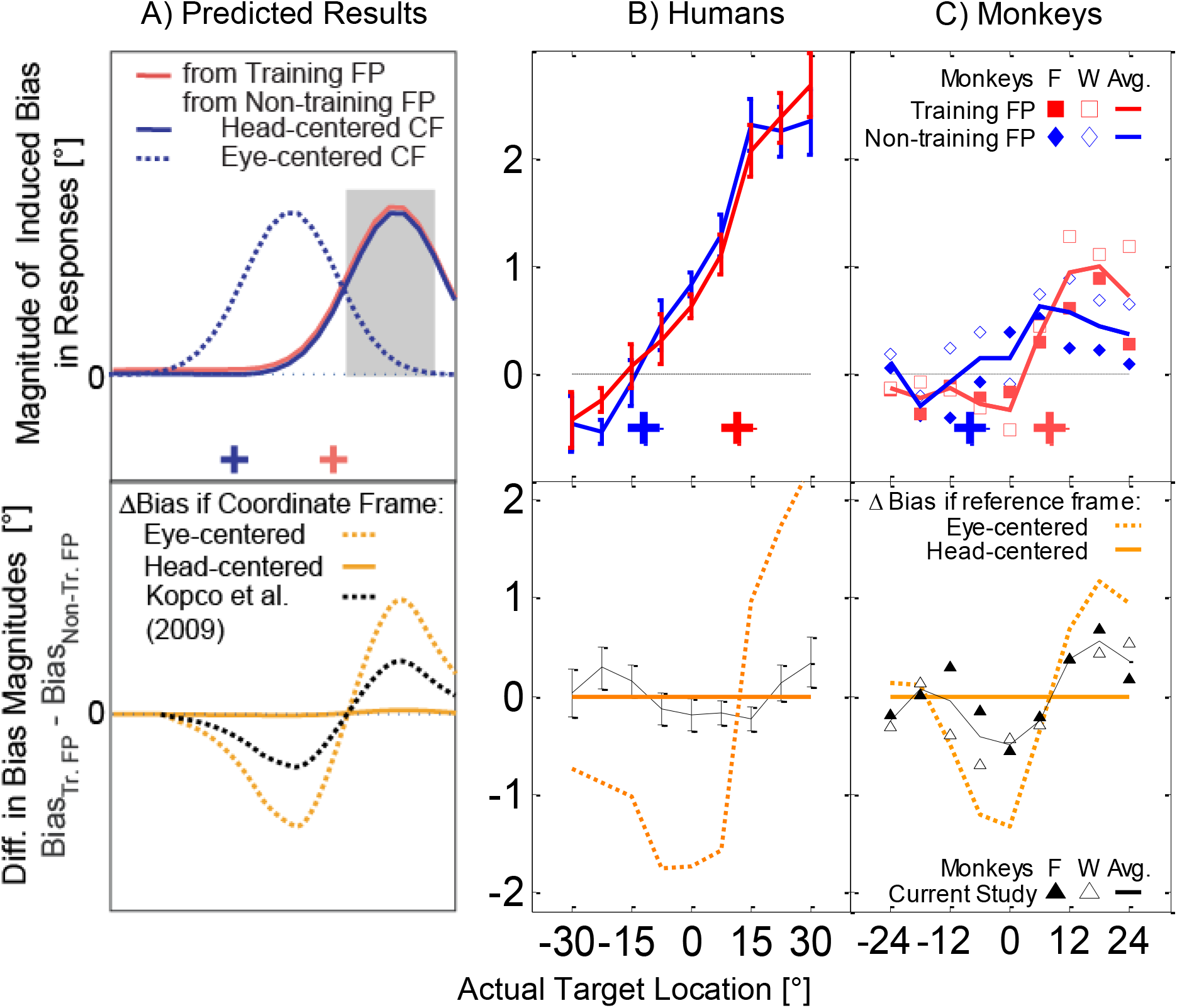
Predicted and observed ventriloquism aftereffect. A) Predicted results when preceding AV trials are presented in the gray region of space (see also Fig. 1A). The plus symbols at the bottom show the training fixation point location (red plus) and the non-training fixation point location (blue plus). The top panel plots the expected pattern of biases induced in the A-only probe responses. Red line shows predictions when the eyes fixate the training FP (i.e., the same FP location used during AV training trials). Solid blue line shows expected results from the non-training FP if the RF of adaptation is head-centered, while dashed blue line shows expected results for an eye-centered RF. The bottom panel shows the differences between the expected bias magnitudes from the training versus the non-training FPs in the two RFs in orange (dashed and solid lines for eye-centered and head-centered, respectively). For comparison, the black line sketches the results corresponding to the mixed RF observed after VAE was induced in the central region in Kopco *et al.* (2009). Upper portion of panels B-C: Magnitude of visually induced shifts in auditory saccades. The graphs show the across-human-subject mean ± 1 SEM (panel B) or the across-monkey-subject mean and individual monkey data (panel C) in the difference between the auditory saccade end point locations when interleaved with spatially displaced AV stimuli vs. when interleaved with AV stimuli with no displacement, collapsed across the direction of the AV displacement. Lower portion of panels B-C: The effect of the initial fixation position on the magnitude of the induced shift: the across-subject mean and ± 1 SEM (panel B) or the individual monkey data (panel C) in the differences between the shifts from the training and non-training FPs (i.e., black lines show the differences between the red and blue lines from the corresponding panels directly above). Orange lines show the predictions of the difference for the two reference frames based on the training FP data (red) from the respective panels above.

We assessed the auditory-only responses interleaved with the spatially mis-aligned AV stimuli against these predictions. The red and blue triangles in Fig. 1B show the raw responses in the conditions in which the ventriloquism aftereffect was induced in a leftward direction (leftward-pointing triangles) or rightward direction (rightward-pointing triangles). Overall, in both humans and monkeys, exposure to spatially mismatched AV stimuli resulted in shift in responses to sounds in the direction of the previously presented visual stimuli (compare the corresponding triangles to the respective circles). To allow a detailed analysis of the results comparable with the predictions of Fig. 3A, red lines in Fig. 3B-C plot the magnitude of the bias in responses measured with eyes fixating the trained FP (red plus sign) *re.* no-shift baseline from Fig. 2, as a function of target location and averaged across the two directions of induced shift (note that no main effect or interaction involving the direction factor were significant in the ANOVA analysis, supporting this way of collapsing the data for visualization; Table 1). The effect was strongest for the three right-most targets, i.e., in the trained region, reaching approximately 2.3° (51% of the ventriloquism effect strength) in the humans and approximately 0.9° (22% of the ventriloquism effect) in the monkeys. It was also location-specific, decreasing quickly toward zero outside of the trained region. These results are consistent with the results of Kopco *et al.* (2009), confirming that the VAE can be induced locally, so that it can be used to assess the VAE RF.

**Table 1.** Three-way repeated-measures ANOVAs of the human and monkey data

| Factor | Humans |  |  | Monkeys |  |  |
| --- | --- | --- | --- | --- | --- | --- |
|  | d.f | F | Signif. | d.f | F | Signif. |
| Speaker Location (1 to 9) | 8, 48 | 33.87 | *** | 8, 8 | 4.5 | * |
| A-only Fixation Point (Tr. vs. Non-Tr.) | 1, 6 | 0.99 |  | 1, 1 | 0.05 |  |
| Direction of Induced Shift (L vs. R) | 1, 6 | 0.43 |  | 1, 1 | 0.09 |  |
| AV Fixation Point (L vs. R) | 1, 6 | 0.27 |  | 1, 1 | 8.37 |  |
| Speaker Location X A-only FP | 8, 48 | 0.79 | * | 8, 8 | 8.88 | * |
| Speaker Location X AV FP | 4, 48 | 2.28 |  | 1, 1 | 0.77 |  |
| A-only Fixation Point X AV FP | 1, 6 | 0.42 |  | 1, 1 | 0.2 |  |
| Speaker Location X Direction | 8, 48 | 0.56 |  | 8, 8 | 1.97 |  |
| AV Fixation Point X Direction | 1, 6 | 2.16 |  | 1, 1 | 0.6 |  |
| A Fixation Point X Direction | 1, 6 | 0.1 |  | 1, 1 | 16.91 |  |
| Speaker Loc. X AV FP X A-only FP | 8, 48 | 0.31 |  | 8, 8 | 0.64 |  |
| Speaker Loc. X AV FP X Direction | 8, 48 | 0.52 |  | 8, 8 | 3.84 |  |
| Speaker Loc. X A-only FP X Direction | 8, 48 | 1.69 |  | 8, 8 | 3.4 |  |
| AV FP X A-only FP X Direction | 1, 6 | 0.12 |  | 1, 1 | 0.59 |  |
| Loc. X AV FP X A-only FP X Direct. | 8, 48 | 1.16 |  | 8, 8 | 0.65 |  |

Significance levels are as follows: \* $p < 0.05$ , \*\* $p < 0.01$ , and \*\*\* $p < 0.005$ .

The reference frame of the VAE was examined by shifting the initial FP to a new location and examining how the observed VAE changed. Blue lines in Fig. 3B-C plot the bias in responses measured with eyes fixating the new, non-trained FP (blue plus sign), shifted by approximately 20° to the left from the trained FP. The effect of shifting the FP differed across the species. In humans, there was very little difference in the measured VAE for the two FPs (blue line lies approximately on top of the red line), while in monkeys, the effect became smaller and its peak moved leftward (the blue line reaches the maximum of approximately 0.5° at the 6° target location). Thus, the observed human results are consistent with visual– auditory recalibration occurring in a predominantly auditory, head-centered, coordinate frame, while the monkey results are again not consistent with either purely head-centered or eye-centered coordinate frame, but intermediate between the two.

To compare the current results more directly to the predictions of the two models and to the data of Kopco *et al.* (2009), a difference between the shift magnitudes from the two FPs was computed (bottom of Fig. 3B-C, black traces) and compared with predictions based on the two models (orange traces). Again, the human results are very close to the predictions of the head-centered RF, while the monkey results fall between the predictions of the two models, suggesting that both the head- and eye-centered signals contribute to visual calibration of auditory space, resulting in a mixed reference frame.

These results were confirmed by performing two separate three-way repeated-measures ANOVAs (one on the human and one on the monkey A-only data), with the factors of target speaker location (nine levels), fixation point of the trials (training vs. non-training FP), and the direction of induced shift (left vs. right). The results of this analysis, summarized in Table 1, show that the main effect of location was always significant, confirming that the ventriloquism aftereffect is spatially specific and does not automatically generalize to the whole audiovisual field. The location by FP interaction was also significant in both species, showing that the reference frame of visual–auditory recalibration is not purely head-centered in either species, even though for humans the eye-centered modulation is relatively small.

## 4. Discussion and Conclusions

The current study examined the spatial properties of the ventriloquism aftereffect induced by AV stimuli presented in only one spatial hemifield in the peripheral audio-visual field. The goal was to ascertain how the ventriloquism aftereffect unfolds as a function of multiple different spatial attributes: fixation position, generalization in head- vs. eye-centered coordinates, and training within one spatial hemifield in contrast to training in both hemifields (as in Kopco et al., 2009). The results indicate that the ventriloquism aftereffect is a multifaceted process, dependent on both the format of the neural representation of space in hearing vs. vision, and on the reference frame used by the two senses.

In terms of the representational format, the location of the fixation position impacted the pattern of adaptation induced by the AV stimuli, even when the AV-stimuli were presented from matching locations and no VAE was induced. This unexpected adaptation was stronger in humans than monkeys, and it was not observed in the previous central-adaptation study (Kopco *et al.*, 2009). It is difficult to identify the cause of this adaptation based on the current data. E.g., since a baseline measurement with no AV stimulation was not performed, it is even hard to tell whether the adaptation, clearly visible in the training-FP – non-training-FP difference plot, is mostly driven by the training-FP or the non-training-FP adaptation. At least two different explanations appear plausible. First, the effect might be a result of adaptation to the auditory stimulus-distribution, which becomes skewed when the training stimuli are included since all of them come from one side (e.g., similar to adaptation reported by Dahmen *et al.*, 2010). Second, the visual signal might be causing some global ventriloquism-like adaptation outside the training region, such that the auditory-only responses are shifted towards the region from which the visual stimuli are frequently presented, but only when the FP is in the hemifield contralateral to the AV stimulation. Whatever the specific mechanism, this adaptation effect shows that there is a hemifield-specific integration of visual and auditory spatial signals that differs from the integration occurring when the stimuli are presented centrally, covering both spatial hemifields.

Regarding the reference frames, the current results together with those of Kopco et al. (2009) show that in humans the RF of VAE is a mixture of eye-centered and head-centered coding in the central region, but mostly head-centered, independent of the direction of the eye gaze, in the periphery. This shows that the transformation of the visual and auditory signals into an aligned reference frame, thought to be necessary for the ventriloquism aftereffect to work, is non-uniform. While it is not immediately clear what form of non-uniformity might be causing this pattern of results, it may be related to the hemispheric-difference channel models of auditory space representation (Salminen *et al.*, 2009; Grothe *et al.*, 2010; Groh, 2014). In monkeys, however, there was no difference between the central and peripheral region data, and the RF was always mixed. Overall, these differences across species and across training regions suggests that the locations in the brain that are recruited to accomplish this recalibration of auditory space may be widely varied. Some are likely head-centered, some are eye-centered, some may involve the position of the eyes in the orbits per se. These sites of plasticity may be recruited differently depending on the training region and whether it spans both head-centered hemifields or is contained within one.

Additional experimental and/or modeling studies are needed to test alternative explanations about the different reference frames of the ventriloquism aftereffect as well as about the unexpected adaptation effect. However, the current results demonstrate that there are hemisphere-specific adaptation processes in visual recalibration of auditory space, resulting in different FP-dependent patterns of adaptation depending on the region in which adaptation is induced.

## Acknowledgments

This work was supported by the SRDA, project DS-2016-0026, EU H2020-MSCA-RISE-2015 grant no. 69122, and by the EU RDP projects TECHNICOM I, ITMS: 26220220182, and TECHNICOM II, ITMS2014+:313011D23. Barbara Shinn-Cunningham was supported by NIH R01 DC013825 Jennifer Groh was supported by NIH NS50942 and NSF 0415634.

